# *Diacamma* ants adjust liquid foraging strategies in response to biophysical constraints

**DOI:** 10.1101/2022.09.13.507744

**Authors:** Haruna Fujioka, Manon Marchand, Adria C. LeBoeuf

## Abstract

Ant foragers need to provide food to the rest of the colony, which often requires transport over long distances. Foraging for liquid is especially challenging because it is difficult to transport and share. Many social insects store liquid food inside the crop to transport it to the nest, and then regurgitate this fluid to distribute it to nestmates through a behaviour called trophallaxis. Some ants instead transport fluids with a riskier behaviour called pseudotrophallaxis – holding a drop of liquid between the mandibles through surface tension. Ants can then share this droplet with nestmates without ingestion or regurgitation. Here, we hypothesised that ants optimise their liquid-collection approach depending on viscosity. Working with a ponerine ant that uses both trophallaxis and pseudotrophallaxis, we investigated why each liquid-collection behaviour might be favoured under different conditions by measuring biophysical properties, collection time, and reaction to food quality for typical and viscosity-altered sucrose solutions. We found that ants can collect more liquid food per unit time by mandibular grabbing than by drinking. At high viscosities, which in nature correspond to high sugar concentrations, ants switched their liquid collection method to mandibular grabbing in response to viscosity, and not to sweetness. Our results demonstrate that ants change their transport and sharing methods according to viscosity – a proxy for sugar concentration in nature – increasing the mass of sugar returned to the nest per trip.

## Introduction

Efficient foraging is crucial for animals to survive, grow and reproduce. Organisms need to balance energy spent with energy gained [1,2]. Optimal foraging theory assumes that animals’ foraging decision making has evolved to the point that the fitness of individuals has been maximised. Foraging provides energy to survive and reproduce; however, it has the cost of exposing the individual to being preyed upon by other animals and it costs energy, for example, due to the time spent exploiting, processing and transporting food.

Animals have a wide variety of foraging strategies. Ants are central-place foragers with diverse diets ranging from complete herbivory to complete predation[3–6]. Morphological and phylogenetic evidence suggests that the ancestral ant was a predator, and transitions to herbivory occurred several times in predatory lineages [5,7,8]. Plant-based food sources, such as plant nectar [9] or honeydew excreted by sapsucking Hemiptera and scale insects [10–12], are rich in carbohydrates. In many species and especially in ecologically dominant ant lineages, these sugary liquids are ants’ main source of energy [13]. Additionally, liquid resources, such as honeydew or nectar, are less ephemeral relative to insect prey and incur fewer risks for ant foragers relative to hunting. Thus, the use of plant-based food sources may lead to lower foraging time and less risk for foragers per calorie returned to the nest.

Transportation of liquid food is a foraging challenge. Many ants and bees transport liquid stored inside their crop, where it cannot easily be lost or stolen [14] during transport. Foragers regurgitate this fluid to distribute it to nestmates through a behaviour called trophallaxis. Many liquid-feeding ants have acquired morphological adaptations for trophallaxis. The ant crop is separated from their midgut by a variably developed proventriculus, which allows the crop to store a large amount of liquid in some species [15]. The structure of proventriculus varies considerably across taxa [16–18], and liquid-feeding ants often have a more elaborate proventriculus. The gaster and crop also need to be expandable to best store liquid food, either temporarily for transport or over the long term in the case of repletes. The extreme example of morphological specialisation are honeypot ants *Myrmecocystus* (Formicinae), where replete workers have a massive ball-like distended gaster full of food to the point where they can barely move [19]. Such species rely on trophallaxis to redistribute food from the repletes to the rest of the colony. Although trophallaxis is considered a safe and reliable liquid transportation method for ants, the crop load (i.e., liquid food intake) strongly depends on these morphological constraints.

Some ants do not have these morphological specialisations but nonetheless consume liquid food: Ectatomminae (*Ectatomma*), Ponerinae (*Diacamma, Neoponera, Odontomachus, Paraponera, Pachycondyla, Rhytidoponera*). These ants typically use mandibular pseudotrophallaxis (hereafter called pseudotrophallaxis) as their method of liquid transport [15,20]. Instead of storing liquid inside the crop, foragers hold liquid food between their mandibles where it forms a droplet maintaining its round shape through surface tension. After foragers return to the nest, they pass the liquid food to nestmates without regurgitation. Previous studies in ponerine ants have reported how this behaviour allows liquids to be distributed in the nest [21,22]. This liquid transport method is sometimes referred to as the ‘social bucket’ method and has been suggested to be an evolutionary precursor to ‘true’ trophallaxis [7].

Handling time is crucial for efficient foraging. For liquid food, the handling time includes both the speed of food collection (i.e., drinking time or grabbing time) and the transport time to the nest. Drinking time in ants has been shown to depend on food quality such as sugar concentration and viscosity itself [23]. Previous studies found that drinking time increased linearly with increasing sugar concentration [23–27]. Individuals need to decide when to stop drinking, considering the balance between energy gain and predation risk. The handling time of pseudotrophallaxis has not been investigated. Regarding transport success, once foragers store food in the crop, they can transport the liquid food safely back to the nest. When using pseudotrophallaxis, there is the possibility of losing the liquid food along the return path. Also, keeping mandibles open adequately may increase the likelihood of predation.

Some ants use both behaviours. The ponerine ant *Diacamma* cf. *indicum* from Japan performs both trophallaxis and pseudotrophallaxis [28], and has a simple proventricular morphology and a rigid, non-extensible gaster. Thus, *Diacamma* cf. *indicum* is an ideal model species to investigate efficient foraging strategies regarding liquid food because it allows us to investigate foraging strategies without morphological specialisation.

The aim of the present study is to reveal what leads ants to use the collection mode of mandibular grabbing instead of drinking and whether ants’ liquid collection modes mechanisms maximise calorie intake rates per foraging trip. We hypothesise that viscosity triggers a switch in collection behaviour between drinking and mandibular grabbing, where mandibular grabbing is more efficient to collect high viscosity solutions. To test this hypothesis, we conducted experiments to investigate 1) physical properties of the liquids such as capillary length, viscosity and surface tension, 2) volume and speed of liquid food collection depending on sugar concentration, 3) whether ants change their transportation method depending on sugar concentration or viscosity by dissociating viscosity from sugar concentration, and 4) the foraging efficiency for each approach using walking speed and the success rate of mandibular transport to estimate the total amount of sugar carried per trip.

## Methods

### Colony collection and colony keeping

Colonies of *Diacamma* cf. *indicum* from Japan were collected from Kenmin-no-mori (Onna) and Sueyoshi park (Naha), Okinawa, Japan. The colonies were kept in plastic artificial nests filled with moistened plaster (9 cm diameter □ × □ 1.5 cm height). Each colony contained a mated gemma-possessing female (i.e., functional queen or gamergate), 50–150 workers, and brood. The artificial nests (90 mm in diameter) were placed in a plastic arena (diameter: cm, height: cm). Nests were maintained at 25 °C under a 12h/12h light-dark regime (light phase: 0800–2000 hours). Reared colonies were fed with chopped frozen crickets three times per week. Water and 10% sugar water were provided *ad libitum*. All behavioural experiments were conducted between 12:00-19:00 in well-lit conditions at 25 □ and 50-60% humidity.

### Behaviour Definitions

The ethogram of the social bucket method (encompassing mandibular grabbing, transport and pseudotrophallaxis) of *Diacamma* ant is shown in Figure 1. Based on a previous study [22] two behaviours for liquid feeding and collection were defined 1) drinking: individuals drink (i.e., mouthpart, labrum, attached to a liquid solution), 2) grabbing: individuals open mandibles to grab and pull at a liquid solution, occasionally succeeding in collecting a droplet.

**Figure 1.**
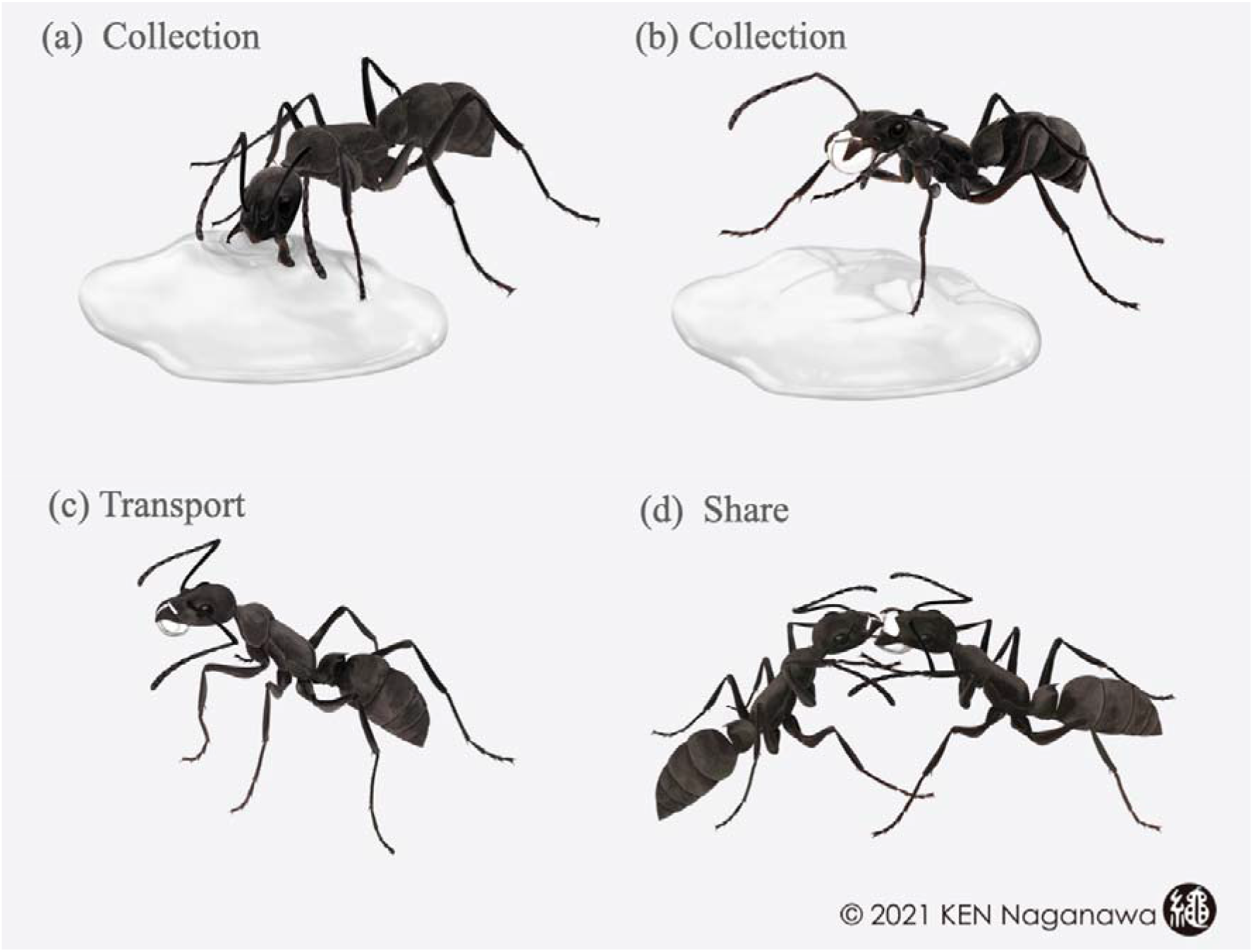
Ethogram of the social bucket technique. An ant touches a solution with the antennae and mouthparts (tasting). After tasting, an individual opens mandibles to grab and pull at the liquid (a). The ant occasionally succeeds in collecting a droplet of solution between the mandibles (b). The individual returns to the nest (c). Inside the nest, the ant shares the drop with other nestmates (d). When the receiver begs for the droplet, their antennae move quickly. Several nestmates can drink from the droplet at the same time. Illustrations by Ken Naganawa.

### Dynamic viscosity measurement

We prepared six sugar solutions, 10, 20, 30, 40, 50, 60 % w/w. The concentrations we used were within the range reported for natural nectar sources (extrafloral nectars: 4.7–76% w/w [29]). In addition, we used two viscosity-altered solutions using carboxymethylcellulose sodium salt (CMC) (Medium viscosity, Sigma-Aldrich), 10% sugar solution with 0.25% CMC (10CMC), and 30% sugar solution with 0.25% CMC (30CMC). The CMC is a non-toxic inert viscosity modifier [30]. By adding this product, we increase the viscosity of the solution without changing its sugar concentration. We confirmed that the sugar concentration of CMC additive solution was not changed using a saccharimeter (Refractometer RBR32-ATC).

We measured these solutions’ dynamic viscosity in a commercial stress-controlled rheometer (ANTON PAAR MCR 300) with a plate-plate geometry of 5 cm diameter and 0.5 mm gap. The bottom plate was roughened by sandblasting to prevent slip artefacts and the temperature was fixed at 25°C by a Peltier hood. We applied constant shear rates, and stress was computed when the steady-state regime had been reached for each shear rate. The resulting stress versus shear rate experiments exhibited linear behaviour characteristic of a Newtonian fluid. This was expected for sucrose/water solutions since sucrose is a small molecule that dissolves well in water, though this was not obvious for the CMC polymer solutions [31]. The dynamic viscosity was directly read from the slope evaluated by least-square minimisation for each sample, for more detail see [32].

### Measurement of surface tension and capillary length

Surface tension is the energy it costs to create a surface of a liquid. We measured the surface tension of the different sugar/water concentrations with a commercial Du Noüy apparatus (Kibron EZPIPlus). The laboratory room was regulated at 25°C +/- 1°C. Each measurement was repeated four times and the reported error is the standard deviation.

Capillary length is the size above which a drop of liquid can no longer sustain its own shape by surface tension but starts to be deformed and to flow under its own weight. In order to obtain the capillary length, we used tabulated density measurements [33]. The capillary length is computed from surface tension and density through the following formula

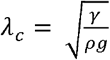

With *γ* the surface tension, *ρ* the density and g the gravitational acceleration [34].

### Drinking speed

Drinking speed was calculated as the slope (μL/sec) of the linear regression of *crop load* and corresponding *drinking time* as intake rate [35]. Thus, we measured these two values (crop load and drinking time) for drinking events of 118 ants from six colonies, drinking seven different solutions: 10, 20, 30, 40, 50, 60% sugar w/w, and 10CMC. The crop load (volume of sugar water drunk) was estimated as the difference in body mass before and after drinking (using a Fisherbrand analytical balance, to 0.1 mg). We chose ants outside of the nest and measured their body mass (before drinking). Then, each individual was separately placed into a plastic box (4.5 □ × □ 4.5 □ × □2 cm), containing a drop (approximately 100 μl) of sugar water. To measure the corresponding drinking time, we video-recorded the behaviour of ants when drinking. Ants were offered one of seven different solutions: 10, 20, 30, 40, 50, 60% sugar w/w, and 10CMC. When ants stopped drinking, we wiped the liquid off the mouthpart with a tissue and measured the ant’s body mass to determine the volume of liquid food inside its crop. We estimated the volume (μL) of sugar water using the average weight (density) of the relevant sugar solution per 1 μl (Supp. table 1). We weighed 100 μL droplets using a micropipette and an electronic balance (Semi-Micro Analytical Balances, GH-202, A&D Company).

We measured the time spent in contact with the droplet by manual observation as the corresponding drinking time. The drinking time was the total time the ant’s mouthpart was attached to a liquid solution. We plotted the crop load and corresponding drinking time and calculated the slope (μL/sec) of the linear regression as the drinking speed (Supp. figure S2). Individuals were never tested more than once a day.

### Measurement of volume of sugar water internally and externally carried

We measured the volume of water internally (crop load) and externally carried (mandible droplet), collected by 207 ants from 4 colonies. We measure the crop load, using the same procedure as ‘drinking speed’. Ants were offered one of six different sugar concentrations: 10, 20, 30, 40, 50, 60% w/w.

Colonies were starved for 3-4 hours before experiments (starvation time based on preliminary behavioural observations, data not shown). The volume of sugar water carried by mandibular grabbing was measured with a microcapillary tube. We placed an artificial nest on one side and a plate-shaped feeder (40 × 40 mm) on the other side of the foraging arena (460 □ × 260 × 100 mm) (Supp. figure 1). Ants were offered one of six different sugar concentrations: 10, 20, 30, 40, 50, 60 % w/w. The ants freely accessed the offered food. After an ant succeeded in grabbing a droplet of the solution, we collected the droplet with a microcapillary tube during a return trip to the nest. The weight of the dispensed volume was measured to calculate the volume of sugar water the ant carried. We estimated the volume (μL) of sugar water using the average weight of the relevant sugar water (Supp. table 1).

### Grabbing time, drinking time and foraging actions in the foraging arena

We measured accumulated grabbing/drinking time during 1047 foraging events of 572 ants from nine colonies. To test the effects of sweetness and viscosity on the foraging methods used, we used seven different sugar concentrations: 10, 20, 30, 40, 50, 60 % w/w, and 10CMC as a viscosity-altered solution. When foragers found 10CMC, they started tasting and drinking as usual. Refusal or avoidance-like behaviours were not observed toward 10CMC.

We placed an artificial nest on one side and a plate-shaped feeder (40 × 40 mm) on the other side of the foraging arena (460 □ × 260 × 100 mm) (Supp. figure 1). Ants were offered one of the seven sugar concentrations: 10, 20, 30, 40, 50, 60 % w/w, and 10CMC. We video-recorded the area around the sugar water droplet for 1 hour. We manually recorded one type of foraging action that foragers had with the droplet, ‘only drinking,’ ‘grabbing after drinking (both),’ or ‘only grabbing.’ Accumulated grabbing/drinking time for each foraging trip was calculated by an observer analysing the videos. The drinking time (sec) is the accumulated time an ant’s mouthparts were attached to the solution. We defined that the start of grabbing was when they opened the mandibles because their mandibles are closed when this focal ant species drinks. The endpoint was when the ant succeeded in grabbing the droplet (e.g., detached droplet from solution). When ants repeated the grabbing, we included grabbing time until the last successful attempt.

### Walking speed and success rate of mandibular transport

We measured ants’ walking speed and success rate using each of the modes of liquid transport. We placed an artificial nest on one side and a plate-shaped feeder (40 × 40 mm) on the other side of the foraging arena (387 × 267 × 65 mm). Ants were offered one of the two sugar concentrations: 10% w/w (low sugar/low viscosity) and 50 % w/w (high sugar/high viscosity). We continuously recorded the foraging arena for 1 hour. The images were captured from above using a web camera (Logicool; HD Pro Webcam C920t) at 10 frames per second. We automatically extracted the coordinates of each individual using the video-tracking system, UMATracker [36]. We categorised ants’ behaviours into 1) empty walking speed, 2) crop-full walking speed, and 3) crop- and mandible-full walking speed by visual observation. We obtained walking speeds for 80 foraging trips from four colonies.

To investigate the success rate of mandibular liquid transport, we observed 30 foraging trips from the feeder to the nest from three colonies and investigated whether ants lost the mandibular droplet during the return trip. The size of the foraging arena (387 × 267 × 65 mm) and two types of sugar water (10% and 50%) were used. If an ant entered the nest without dropping the mandibular droplet, we recorded that as a foraging success. Returning to the nest after drinking was always considered a foraging success as the liquid cannot be lost.

### Estimation of sugar intake

To assess foraging efficiency, we estimated total sugar intake per foraging trip by combining several forms of data: 1) observation of foraging action ants performed, 2) drinking or grabbing time, 3) drinking speed, and 4) average grabbing volume. First, we calculated the total liquid load per trip based on the foraging action used (see the section *‘Grabbing time, drinking time and foraging actions in the foraging arena’* and Figure 4). The total crop load per trip was estimated by multiplying the drinking speed (see the section *‘Drinking speed’*, Supp. figure 2) by the accumulated time drinking in the foraging arena (see the section *‘Grabbing time, drinking time and foraging actions in the foraging arena’*, Figure 3b). The mandible load (collected by grabbing) was defined as the average volume of liquid carried for each sugar concentration in the experiment *‘Measurement of volume of sugar water internally and externally carried’* (see Figure 3a). When ants performed both drinking and grabbing, we summed the average volume of liquid carried and the estimated total crop load. Using the average weight of sugar water (Supp. table 1), we converted the total liquid load per trip (μL) to the weight of the liquid load (mg). From the liquid load weight (mg), we calculated the total sugar intake per trip (mg) for each sugar concentration.

### Statistical analysis

Linear regression models were used to calculate the drinking speed. We compared regression slopes using a Wilcoxon test with Bonferroni correction. Generalised linear regression models were used to investigate the relationship between the liquid volume or loading speed with the food quality variables and foraging actions. Pairwise chi-square tests with a Bonferroni correction were used for comparing foraging action on different sugar concentrations. The relationship of the load with the food quality variables and foraging actions was analysed by two-way analysis of variance (ANOVA), followed by Tukey’s honestly significant difference (HSD) test for multiple comparison of means. A significance level of 5% was used in all comparisons. All analyses were run in R studio 2022.02.3 (package: ggplot2), except for viscosity measurements and drinking speed that were analysed in python 3.9 and the linear regressions were done with the module scipy.optimize v1.7.1. [37].

## Results

*Diacamma* ants forage for sugary liquids and use two forms of behaviour to collect liquid – drinking and mandibular grabbing. To understand whether ants alter their liquid collection and transport behaviour according to the contents and properties of the liquid they are transporting, we measured multiple variables to find what ants are optimising: volume carried, time spent collecting, frequencies of different foraging actions, walking speed, success rate of transport, biophysical properties of the liquids and sugar load acquired per trip.

First, we analysed three liquid properties of a range of ecologically relevant sugary liquids from 0-60% sucrose: dynamic viscosity, surface tension, and capillary length. Dynamic viscosity increased with increasing sugar water concentration (Figure 2a). We altered the viscosity of a low-sugar solution using the viscosity-modifying additive CMC (carboxymethylcellulose sodium salt). The viscosity of 10% sugar water with CMC (10CMC) was comparable to the viscosity of 40-50% sugar water (Supp. Table 2, Figure 2a). We measured an increase in surface tension (mN/m) with increasing sugar concentration while the capillary length was not significantly affected (Table 1).

**Figure 2.**
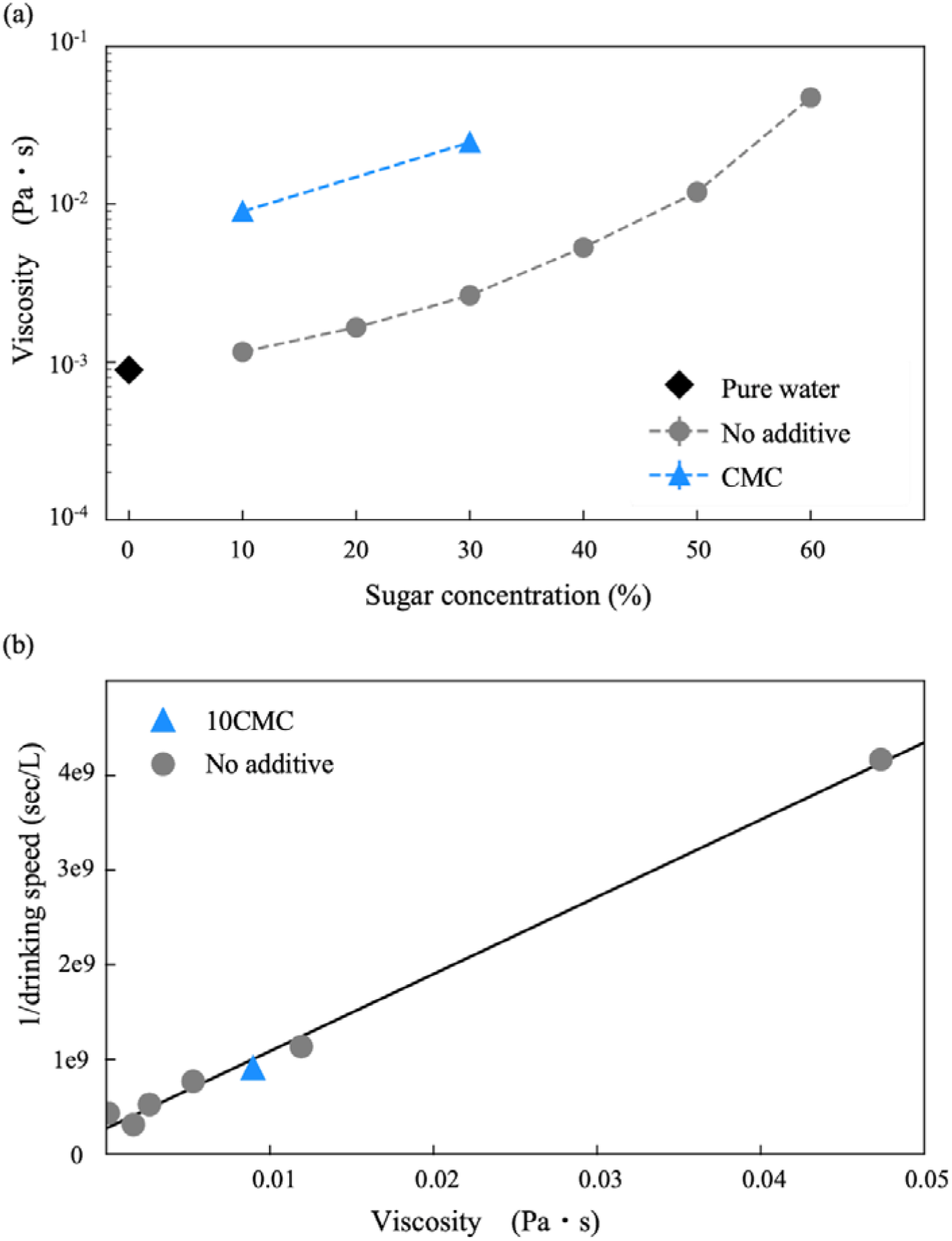
Dynamic viscosity leads to drinking speed. (a) Dynamic viscosities at 25°C for different sugar water solutions as a function of the sugar concentration (w/w). The different colours correspond to pure water (circle) and viscosity-altered solutions (triangle). The error bars denote the quality of the Newtonian fit applied to each flow curve for each solution. The blue triangle indicates the viscosity-altered solutions 10CMC and 30CMC sugar solution with 0.25 CMC(w/w). (b) The relationship between drinking speed and viscosity. The inverse of drinking speed increases linearly with dynamic viscosity for sugar solutions with or without viscosity-altering additives.

**Figure 3.**
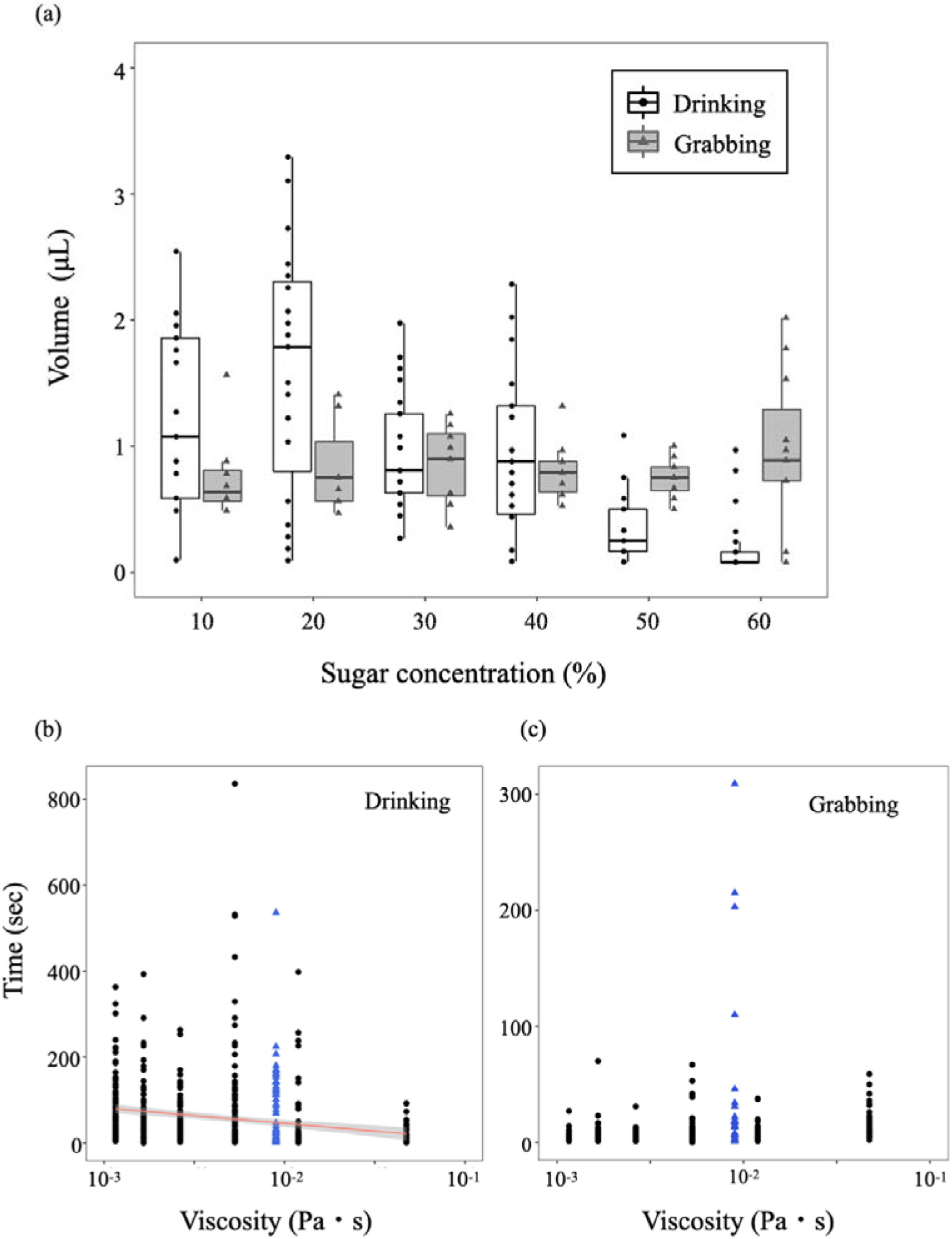
Impact of sugar concentration and viscosity on liquid collection. (a) The amount of liquid collected by drinking (black) and grabbing (grey) at different concentrations of sugar water (% w/w). Time spent drinking (b) and grabbing (c) by ants at different concentrations and viscosities of sugar water (% w/w). Six sugar concentration solutions are shown in black circles, and the viscosity-altered solution 10CMC is shown with blue triangles. Statistical analysis can be found in Table 2. Linear regression shown in (b) is performed only on the unaltered solutions…

**Figure 4.**
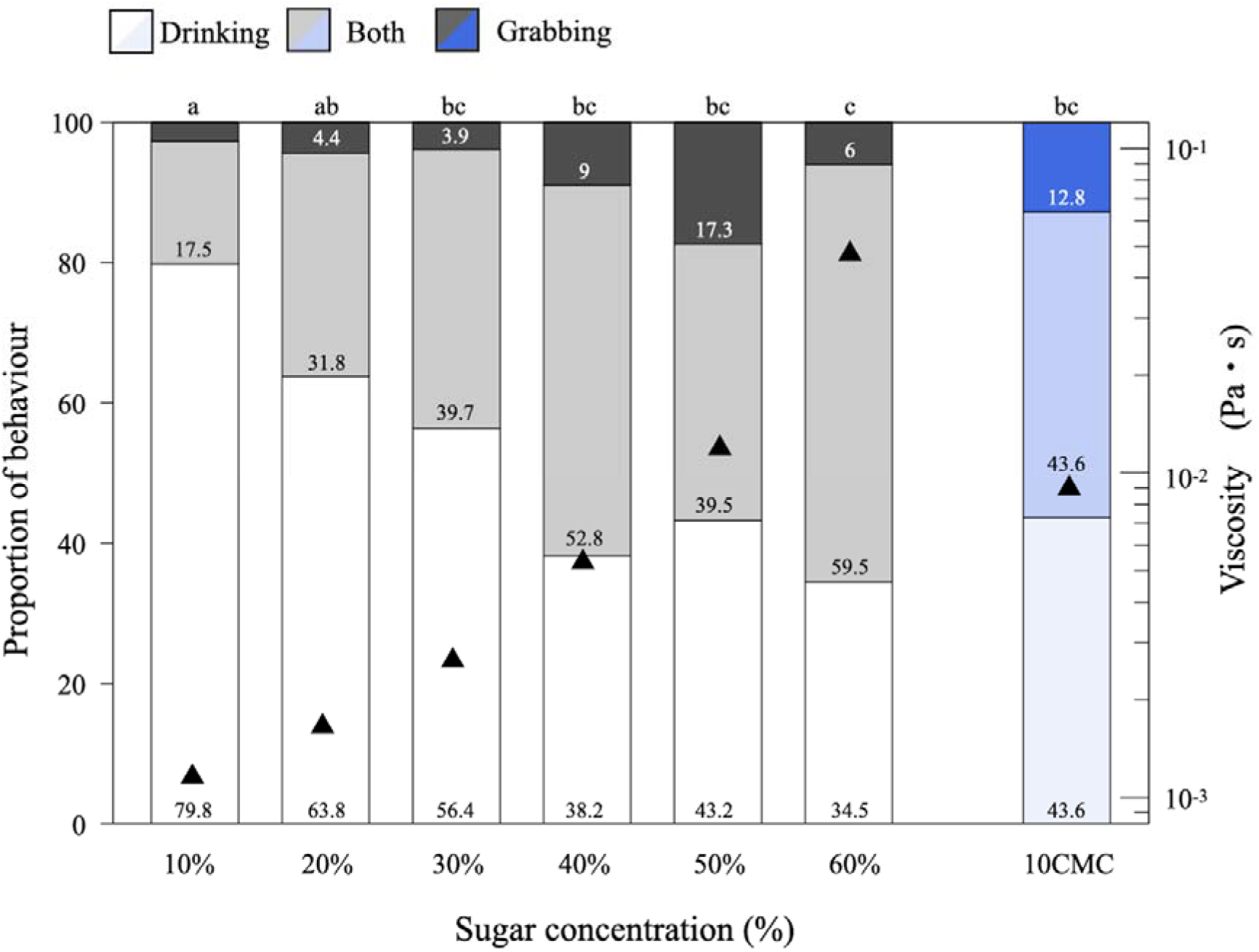
The proportion of pseudotrophallaxis depends on sugar concentration and viscosity. Drinking (white), both (grey), and grabbing (dark grey) indicate the behaviour of only pseudotrophallaxis, pseudotrophallaxis after drinking, and only drinking. The blue-ish bars indicate the viscosity-altered solution 10CMC. Letters on the top of the bar mean they were significantly different at P < 0.05 (chi-sq test with Bonferroni correction).

**Table 1.** Surface tension and capillary length for water/sugar solutions.

| Sugar<br>concentration | Surface tension<br>(mN/m) | Std. Error<br>surface<br>tension<br>(mN/m) | Density<br>(g/mL) | Capillary<br>length<br>(mm) |
| --- | --- | --- | --- | --- |
| 0 | 71.7 | 0.1 | 0.9982 | 2.71 |
| 10 | 70.7 | 0.5 | 1.022 | 2.66 |
| 20 | 74.7 | 0.4 | 1.0631 | 2.68 |
| 30 | 80.0 | 0.4 | 1.1132 | 2.70 |
| 40 | 77.8 | 0.3 | 1.1374 | 2.64 |
| 50 | 78.8 | 0.3 | 1.1975 | 2.59 |
| 60 | 84.5 | 0.3 | 1.2391 | 2.64 |

Combining video recording and weight measurements of ants before and after drinking, we determined that drinking speed is inversely proportional to the viscosity of the sugar water solution imbibed (Supp. Table 3, Figure 2b). If we approximate flow as if the alimentary canals of ants were cylindrical pipes, we can use the Hagen-Poiseuille equation

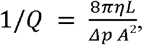

where Q is the drinking speed, L and A the length and cross-sectional area of the ant throats, *η* the viscosity of the solutions and Δ*p* the pressure drop that the ants generate by suction [38]. Thus, high viscosity makes drinking more time-consuming and may cause the ants to switch toward grabbing behaviour. Indeed, drinking speed for 10CMC was slower than that for 10% sugar water (Wilcoxon test with Bonferroni correction, 10CMC vs. 10%: p = 0.013) and comparable to drinking speed for 40% and 50% sugar water (Wilcoxon test with Bonferroni correction, 10CMC vs. 40: p = 0.58; 10CMC vs. 50: p = 0.37). Drinking speed for 10CMC falls onto the linear adjustment for drinking speed for solutions without additives (Figure 2b).

We investigated the volume of liquid food collected by ants using two different collection methods, feeding on different sugar concentrations (Figure 3). For the volume ants collected through drinking or mandibular grabbing, we observed an interaction between sugar concentration and foraging action (Figure 3a, Table 2a, GLM, sugar × collection method: p < 0.001), and therefore we analysed the foraging actions separately. When ants drank liquid, the amount imbibed decreased as viscosity increased (Figure 3b, Table 2b, LM: p < 0.001). For the amount of liquid grabbed within the mandibles, there was no significant trend across the different viscosity (Figure 3c, Table 2b, LM: p = 0.14).

**Table 2.**
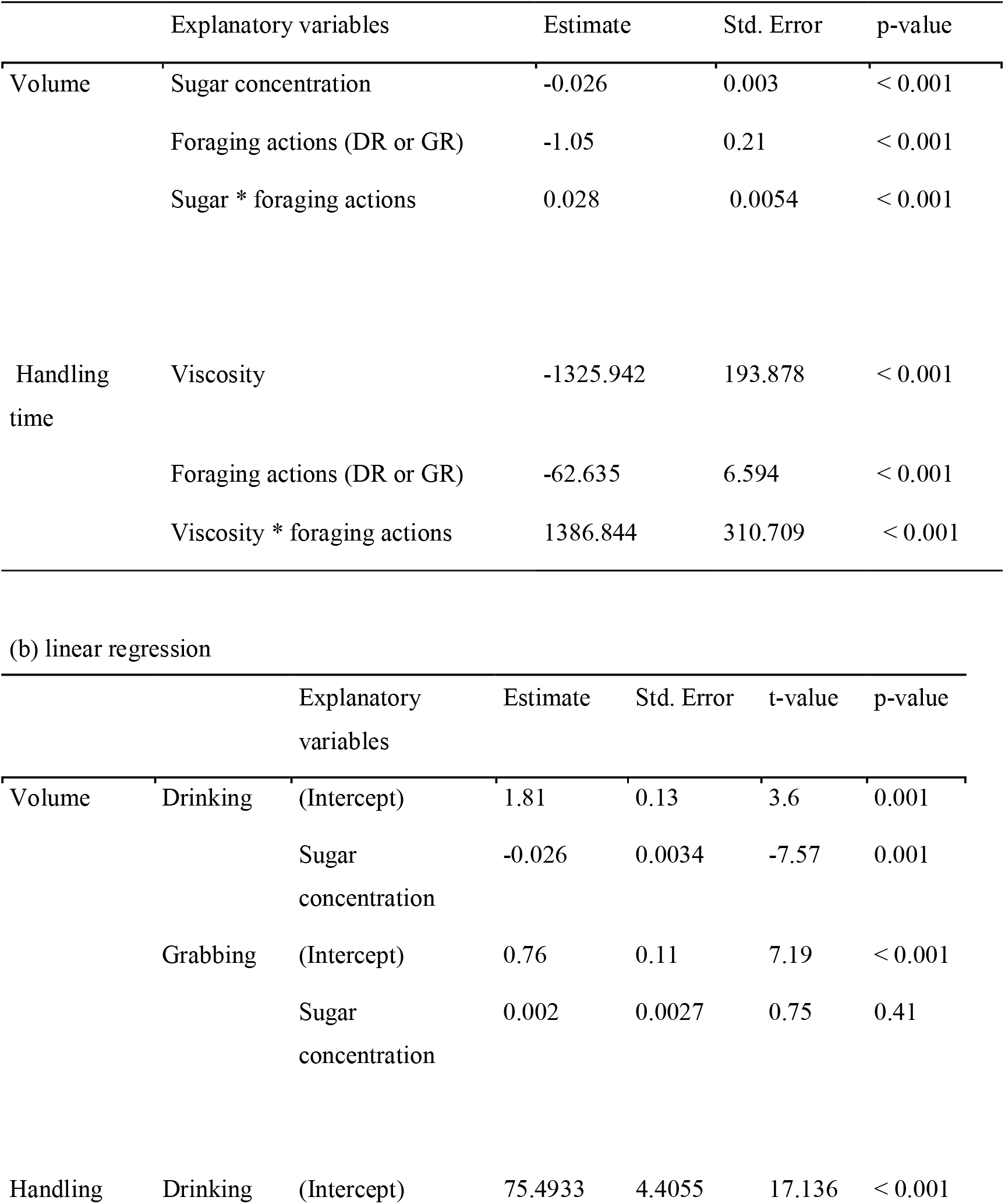
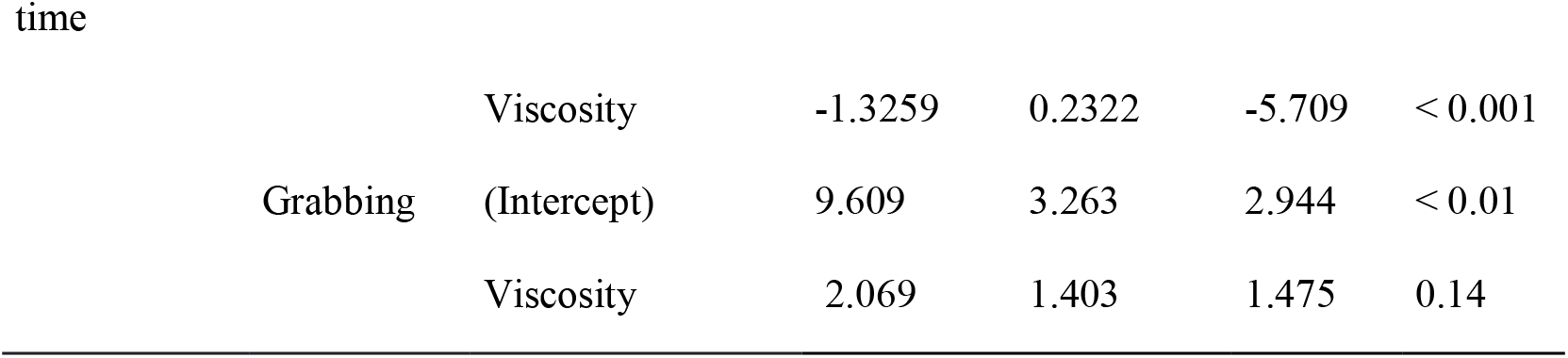
Influence of sugar concentration and foraging action on volume and handling time. Results of generalised and general linear regression (a) and linear regression model (b) between two explanatory variables and volume of carried or drunk water and handling time. Sugar concentration water is 10, 20, 30, 40, 50, and 60% (w/w) and foraging actions are drinking (DR) and mandibular grabbing (GR). Viscosity is seven different sugar and viscosity solutions; 10, 20, 30, 40, 50, 60% (w/w), and the viscosity-altered solution 10CMC.

Since the volume of droplets collected by mandibular grabbing were more or less constant (~1μL) at all sugar concentrations (Fig 3a), the energy required to extract the droplets should be solely a function of the surface tension, where it should be easier to grab lower sugar concentration solutions due to the lower surface tension (Table 1). In terms of capillary length, our measurements showed that the increase in surface tension with the addition of sugar is balanced by the increase in density (Table 1). These values lead to an estimate of the maximum transportable volume as a drop of approximately 80 ± 10 μL, estimated from the capillary length translated into the volume of a sphere. This volume is more than one order of magnitude above the maximum volume measured in this study (Table 1, Figure 3a). Thus, at all sugar concentrations, the droplets transported by grabbing could sustain their own shape, were not impacted by gravity and did not flow during transport. This suggests that the limiting factor of the grabbing volume is not biophysical liquid properties, but more likely the ants’ morphological constraints.

Regarding the collection time for drinking or mandible grabbing, we found an interaction between viscosity and foraging action (Table 2a, GLM, viscosity × foraging action: p < 0.001). When ants drank, the drinking time decreased with increasing viscosity (Figure 3b, Table 2b, LM: p < 0.001). There was no linear relationship between grabbing time and viscosity (Figure 3c, Table 2b, LM: p = 0.14). All in all, mandible grabbing took less time to collect liquid when compared to drinking. Thus, mandible grabbing is a more effective method to collect liquid food, in terms of volume per foraging time, across different sugar concentrations.

In spite of this, ants often collected sugar water in their mandibles after drinking and rarely performed only mandible grabbing without drinking (Figure 4). The proportion of these foraging actions was significantly different across sugar concentrations (Figure 4, p < 0.05, chi-sq test with Bonferroni correction). The proportion of mandible grabbing after drinking (both) and mandible grabbing alone (both of which result in pseudotrophallaxis) increased with increasing sugar concentration. This indicates that ants switch to grabbing and pseudotrophallaxis when they feed on liquid food with higher concentrations of sugar. This could come about because this high-sugar food is more valuable or because high-viscosity liquids are difficult for them to drink, as indicated by the drinking speed measurements (Figure 2). To test whether ants react to changes in sugar concentration or viscosity, we offered ants our viscosity-altered solution 10CMC and observed which foraging action was used. The proportion of ants drinking significantly decreased compared to those drinking 10% sugar water (Figure 4, p < 0.05, chi-sq test with Bonferroni correction). The proportion of ants drinking 10CMC was equivalent to those drinking a high viscosity 50% sugar solution (Figure 4). This result shows that ants switch collection methods in response to viscosity, and not to sweetness.

To investigate whether this transition toward grabbing over drinking with increasing viscosity is related to foraging efficiency, we estimated the liquid load and sugar load per trip across the different sugar concentrations. The liquid load per trip had the largest volume when ants used both drinking and grabbing in the same trip (Figure 5a). There was a significant interaction between sugar concentration and foraging action on the liquid load (Table 3, ANOVA, sugar × foraging action: p < 0.01). The total liquid load when grabbing was larger than the crop load (Table 3, ANOVA, foraging action: p < 0.05). The crop load decreased with increasing sugar concentration (Tukey’s HSD Test, p < 0.001). To examine how much energy ants can bring back to the nest through these methods, we transformed the liquid load to sugar load. We found that the difference in efficiency between drinking and grabbing increased with sugar concentration (Figure 5b). There was a significant interaction between sugar concentration and foraging action (Table 3, ANOVA, sugar × foraging action: p < 0.01). The sugar loads of drinking were higher in the 20-40% of sugar concentrations (Wilcoxon test with Bonferroni correction, p < 0.05). The total sugar load acquired by grabbing significantly increased with sugar concentration (Wilcoxon test with Bonferroni correction, p < 0.05). In addition, for the high viscosity and low sugar water 10CMC, the total sugar load per trip was lower than 10% sugar water (Figure 5b, Wilcox test with Bonferroni correction, p < 0.05). Compared to the solution of similar viscosity (40% sugar water), the sugar intake of 10CMC was less per foraging trip (Wilcoxon test with Bonferroni correction, p < 0.05). If ants were optimising sugar returned to the nest per unit time while collecting (as opposed to per trip), grabbing only would be the optimal collection behaviour across viscosities.

**Figure 5.**
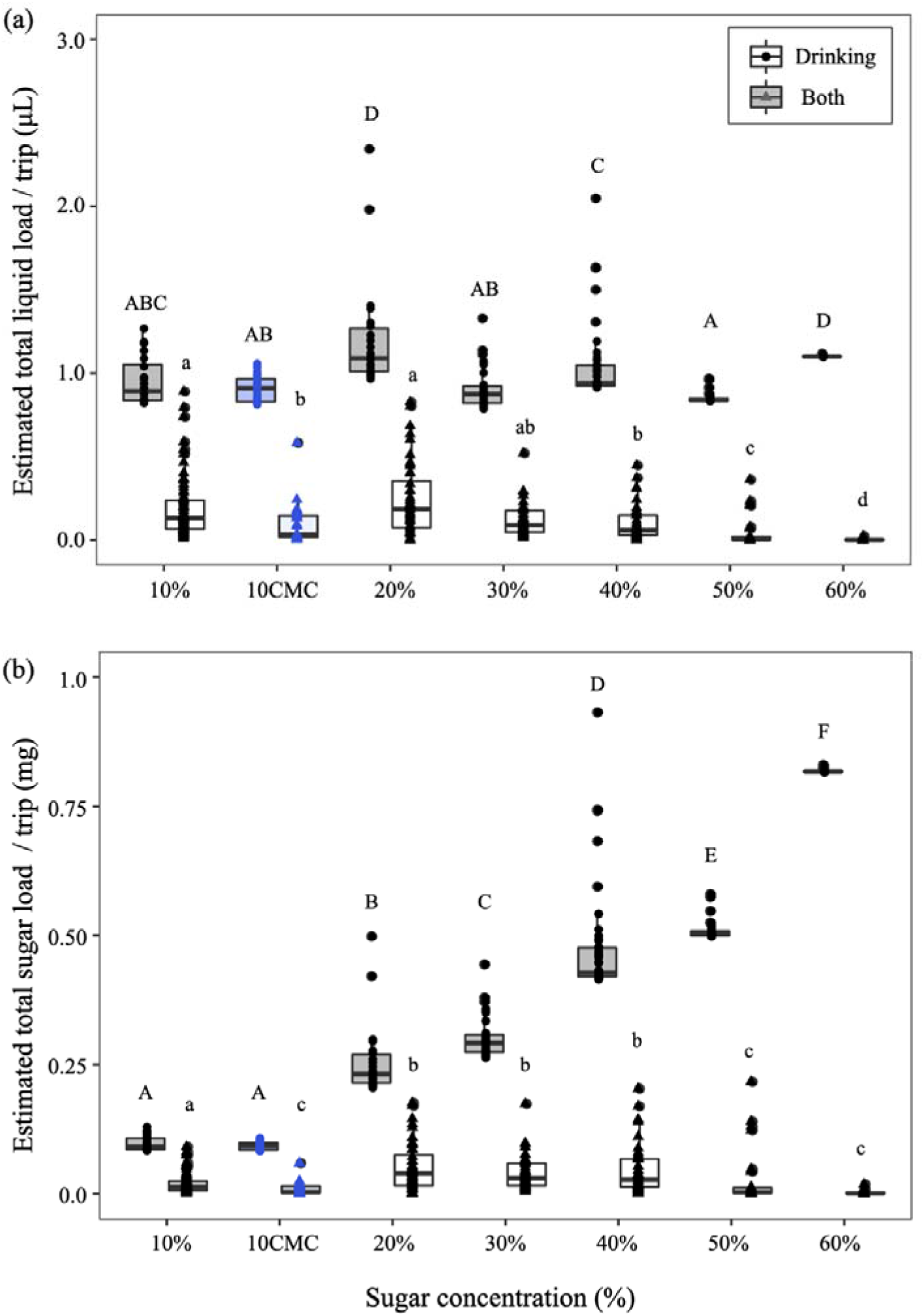
Foraging efficiency for the foraging action. Estimated total liquid (a) and sugar load per trip (b) are estimated based on drinking time and foraging action. White (circle) and dark grey (triangle) boxes indicate the action of drinking and both drinking and grabbing. Blue indicates the 10CMC viscosity-altered solution. Statistical analysis can be found in Table 3. Letters above points indicate significant differences of p < 0.05 (Wilcoxon test with Bonferroni correction).

**Table 3.** Influence of different concentrations and viscosity of sugar water and foraging action on liquid and sugar load. Results of two-way ANOVA between two explanatory variables and liquid load or sugar load per trip. Solutions are seven different sugar solutions 10, 20, 30, 40, 50, and 60% (w/w) and 10CMC. Collection methods are drinking (DR) and both drinking and mandibular grabbing (both).

|  | Explanatory variables | Df | Sum Sq | Mean Sq | F value | p-value |
| --- | --- | --- | --- | --- | --- | --- |
| Liquid load | Solutions | 6 | 8.66 | 1.44 | 60.12 | < 0.001 |
|  | Collection methods (DR or both) | 1 | 92.13 | 92.13 | 3838.72 | < 0.001 |
|  | Solution * collection methods | 6 | 1.55 | 0.26 | 10.74 | < 0.001 |
|  | Residuals | 515 | 12.36 | 0.02 |  |  |
| Sugar load | Solutions | 6 | 13.758 | 2.293 | 1154.6 | < 0.001 |
|  | Collection methods (DR or both) | 1 | 14.902 | 14.902 | 7503.6 | < 0.001 |
|  | Solution * collection methods | 6 | 6.841 | 1.140 | 574.1 | < 0.001 |
|  | Residuals | 515 | 1.023 | 0.002 |  |  |

To understand why ants might drink as well as grab, we investigated how often failures occur during mandibular transportation and whether carrying a mandibular droplet causes ants to walk more slowly. In our experimental set-up, a flat and relative short foraging path (approximately 30 cm), all ants succeeded in transportation by mandibular grabbing (Supp. Table 4). For both mean walking speed and maximum walking speed, there were no significant differences across the three categories: empty-crop, crop-full, and crop- and mandible-full (Supp. Fig. 4, Tukey-Kramer test). Even when full-crop ants held a droplet in their mandibles, their walking speed did not decrease compared to ants in empty-crop and full-crop conditions (Supp. Fig. 4).

## Discussion

Behaviour in a given species is highly adapted to that organism’s context, and these behavioural adaptations often involve precise forms of behavioural plasticity. In this study, we analysed the flexibility of foraging behaviours in response to viscosity in *Diacamma* cf. *indicum*. The ant used two liquid collection actions – drinking and grabbing – when collecting liquid food. In this study, we aimed to quantify dynamic switching between of two types of liquid food collection used by a single ant species. *Diacamma* cf. *indicum* is a part of a clade of ants that rarely specialise on liquid food. Given this species’ phylogenetic context, these behaviours are likely to be relatively recent specialisations [15]. However, it remains unclear when ants use pseudotrophallaxis as opposed to true trophallaxis to collect, transport, and share liquids, whether the use of these behaviours varies according to food quality, and whether one is an evolutionary step toward another.

### Viscosity dictates behaviour

Here, we clearly observed mandibular gabbing and pseudotrophallaxis in the lab in *Diacamma* cf. *indicum* and saw that their use of this collection behaviour changes with sugar concentration and viscosity. Our results are consistent with previous studies in other ponerine ants, where ants stopped drinking [39] and tended to use mandibular grabbing at higher sugar concentrations (> 40%) [40]. By decorrelating viscosity from sugar content, our work revealed that ants made this switch in collection mode according to viscosity only (Figure 4). Viscosity has been seen to reduce the liquid intake rates in many insects, including ants [23,39,41–43], consistent with our results (Figure S2). Here, we directly measured the drinking speed of the ants. We show a robust linear relationship between the inverse of drinking speed and the viscosity of the solutions regardless of the presence of the viscosity-altering agent, indicating that viscosity is the main factor regulating drinking time. Our analyses of surface tension and capillary length indicate that the biophysical limits for maximum droplet volume of these solutions that *Diacamma* ants could transport through surface tension are never reached. Rather, the constraints on volume obtained by grabbing are more likely related to the ants’ size and mandible span, which could be explored in future studies for example in wasps that are physically larger and that also transport fluids between their mandibles.

We found that ants used mandibular grabbing after drinking liquid (Figure 3a). This maximises liquid load per trip because ants can transport both internally and externally. Multiple trips may be costly as they involve loss of energy and increased predation risk. We observed no foraging failures or reduced travel speeds when ants were internally and externally loaded. Thus, the key factor in evaluating foraging efficiency would be calorie intake per trip. The data showed that the total sugar load clearly increased at higher sugar concentrations when ants used mandibular grabbing, suggesting that grabbing, and consequently pseudotrophallaxis, are more efficient methods to collect high-viscosity liquid than drinking and regurgitation.

One possibility is that ants drink to fulfill individual energetic needs, but transport liquid through grabbing for the community. The sugar load by drinking 10% sugar water was greater than the high-viscosity 10CMC solution (Figure 5). That might be caused by the fact that ants can drink less high-viscosity fluid per unit time than low-viscosity fluid. If the volume of liquid imbibed was solely taken to cover individual energetic foraging costs, they would drink the same *sugar content*, which was not the case here.

### Why do they not always use both drinking and mandibular grabbing?

One possibility is that mandibular grabbing is a risky, but high pay-off collection method. For example, if ants encounter predators, they might not react quickly enough, ending up to lose their mandibular droplet and/or being preyed upon because they are less agile. Ants might not use mandibular grabbing in dangerous sites where they encounter predators. On the contrary, if there are competitors around the food site, ants need to compete against other ant species. *Pheidole megacephala* soldiers reacted to the presence of competitors. The soldiers performed more mandibular grabbing on the territory of other ant species in order to rapidly gather and transport large loads of liquid food [44]. Future studies with different distances and ecological contexts while analysing transport time could help elucidate the cost of transportation of pseudotrophallaxis through surface tension.

In addition, the speed of sharing to nestmates may differ between trophallaxis and pseudotrophallaxis. After foragers return to the nest, they share the liquid food through trophallaxis (regurgitation), or through pseudotrophallaxis. Trophallaxis should more rapidly distribute food in the colony because a receiver can become a donor and continue to distribute liquid food by regurgitation, also allowing the formation of a more complex social network [45–47]. In the carpenter ant *Camponotus*, foragers give food to a receiver, proportional to the available capacity in the receiver’s crop. This trophallactic interaction helps the forager to sense colony satiation level and decide when to leave the nest and bring in more food [48]. However, the distribution dynamics of liquid by pseudotrophallaxis have not been studied. The observation of liquid distribution in the focal species that use both trophallaxis and pseudotrophallaxis is needed to understand these dynamics of liquid distribution and the regulation of foraging effort.

### Why do so few ant species perform mandibular grabbing?

Mandibular grabbing and pseudotrophallaxis are mostly performed by ponerine ants (Ponerinae), with only a few noted exceptions in other major ant subfamilies [38] One possible reason why some ants use pseudotrophallaxis and others use true trophallaxis is that many ants have internal morphological adaptations for liquid intake, storage and regurgitation, such as an expandable gaster, an elastic crop, a larger mouth and throat, and a highly developed proventriculus [7,15]. These adaptations possibly allow for greater flexibility regarding the intake of high-viscosity solutions [16,39]. Thus, species with these morphological adaptations may not need pseudotrophallaxis.

A second possibility is that there might be biophysical restrictions on whether an ant can collect a liquid drop between her mandibles. It is likely that small body size makes interactions with liquid droplets more dangerous due to the difficulty of getting out of a liquid with a high surface tension [49]. We observed that ants make a ‘hasty’ motion at the end of the extraction of the droplet. The force needed to break the droplet away is also directly dictated by the surface tension of the liquid. For small ants, it may be more difficult to exert the force required to extract the droplet. Whether any relationship exists between the ability to perform pseudotrophallaxis and biophysical restrictions, related to body size, has not yet been studied, making this area well poised for study from a biomechanics perspective.

### Share with nestmates: socially transferred materials

Trophallaxis allows for medium- to long-term food storage before redistribution while pseudotrophallaxis does not. Thus, ecological contexts and environmental harshness may also tilt an ant to engage in one behaviour or another. Another valuable feature of trophallaxis is that donors can alter the contents of what they pass to nestmates, either through partial digestion or through more complex signalling [50–52], which may bias a species or even a single ant to use one or another behaviour. Recent studies reveal that ants’ regurgitated fluid contained more than food [50,51]. For example, trophallactic fluid in carpenter ants contains hormones, nestmate recognition cues, small RNAs, and various proteins. In *Diacamma*, it is unclear whether foragers regurgitate the contents of their crop during pseudotrophallaxis, or if they only regurgitate when they do trophallaxis. Future studies could examine whether they add any endogenous materials during these behaviours.

## Data accessibility

All data and codes are available at: https://datadryad.org/stash/share/zYykgqIg6GgOgRjhPVqO4Lwq-Xhe8etdBlHl4qHQZhg

## Fudging

This work was funded by JSPS KAKENHI [Grant number JP20J01766] and Myohara Maroko research grant by the Zoological Society of Japan to HF, and Swiss National Science Foundation Grant PR00P3_179776 to ACL.

## Acknowledgments

We are grateful to Ken Naganawa for his amazing illustrations. We would like to thank Dr. François Lavergne and Satomi Koga for their contribution to data collection and Dr. Andrew Brown for his advice on statistical analysis. We also thank all members of the social fluids lab at University of Fribourg for discussions about the study and Dr. Isaac Planas-Sitjà and Marie-Pierre Meurville for providing feedback on early drafts of the manuscript.

## References

1. Pyke GH, Pulliam HR, Charnov EL. 2015 Optimal Foraging: A Selective Review of Theory and Tests. https://doi.org/10.1086/409852 52, 137–154. (doi:10.1086/409852)

2. Stephens DW, Krebs JR. 1987 Foraging Theory. Princeton University Press. (doi:10.1515/9780691206790)

3. Buckley RC. 1987 Interactions involving plants, Homoptera, and ants. Annual review of ecology and systematics. Vol. 18, 111–135. (doi:10.1146/annurev.es.18.110187.000551)

4. Davidson DW, Cook SC, Snelling RR, Chua TH. 2003 Explaining the abundance of ants in lowland tropical rainforest canopies. Science (1979) 300, 969–972. (doi:10.1126/SCIENCE.1082074/SUPPL_FILE/DAVIDSON.D.SOM.PDF)

5. Nelsen MP, Ree RH, Moreau CS. 2018 Ant–plant interactions evolved through increasing interdependence. Proceedings of the National Academy of Sciences 115, 12253–12258. (doi:10.1073/pnas.1719794115)

6. Heil M, Mckey D. 2003 Protective Ant-Plant Interactions as Model Systems in Ecological and Evolutionary Research. Source: Annual Review of Ecology, Evolution, and Systematics 34, 425–453. (doi:10.1146/132410)

7. Hölldobler B, Wilson EO. 1990 The Ants. Cambridge: Harvard University Press.

8. Wilson EO. 1990 The insect societies (Harvard paperbacks).

9. Calixto ES, Lange D, Del-Claro K. 2021 Net benefits of a mutualism: Influence of the quality of extrafloral nectar on the colony fitness of a mutualistic ant. Biotropica 53, 846–856. (doi:10.1111/BTP.12925)

10. Pekas A, Tena A, Aguilar A, Garcia-Marí F. 2011 Spatio-temporal patterns and interactions with honeydew-producing Hemiptera of ants in a Mediterranean citrus orchard. Agric For Entomol 13, 89–97. (doi:10.1111/j.1461-9563.2010.00501.x)

11. Pringle EG. 2021 Ant-Hemiptera Associations. Encyclopedia of Social Insects, 45–48. (doi:10.1007/978-3-030-28102-1_8)

12. Styrsky JD, Eubanks MD. 2006 Ecological consequences of interactions between ants and honeydew-producing insects. Proceedings of the Royal Society B: Biological Sciences 274, 151–164. (doi:10.1098/RSPB.2006.3701)

13. Sempo G, Detrain C. 2004 Between-species differences of behavioural repertoire of castes in the ant genus Pheidole: A methodological artefact? Insectes Soc (doi:10.1007/s00040-003-0704-2)

14. Richard FJ, Dejean A, Lachaud JP. 2004 Sugary food robbing in ants: a case of temporal cleptobiosis. C R Biol 327, 509–517. (doi:10.1016/J.CRVI.2004.03.002)

15. Meurville MP, LeBoeuf AC. 2021 Trophallaxis: the functions and evolution of social fluid exchange in ant colonies (Hymenoptera: Formicidae). Myrmecol News 31, 1–30. (doi:10.25849/MYRMECOL.NEWS_031:001)

16. Davidson DW, Cook SC, Snelling RR. 2004 Liquid-feeding performances of ants (Formicidae): Ecological and evolutionary implications. Oecologia 139, 255–266. (doi:10.1007/S00442-004-1508-4/TABLES/4)

17. Eisner T. 1957 A comparative morphological study of the proventriculus of ants (Hymenoptera: Formicidae). Bull Mus Comp Zool 116, 437–490.

18. Eisner T, Brown WLJ. 1958 The evolution and social significance of the ant proventriculus. Proceedings Tenth International Congress of Entomology 2, 503–508.

19. Robinson EJH, Feinerman O, Franks NR. 2009 Flexible task allocation and the organization of work in ants. Proceedings of the Royal Society B: Biological Sciences 276, 4373–4380. (doi:10.1098/rspb.2009.1244)

20. Hölldobler B. 1985 Liquid food transmission and antennation signals in ponerine ants. Isr J Entomol, 89–99.

21. A. Dejean, J.P. Suzzoni. 1997 Surface Tension Strengths in the Service of a Ponerine Ant: a New Kind of Nectar Transport. Naturwissenschaften 84, 76–79.

22. Lachaud Jean-Paul, Alain Dejean. 1991 Food sharing in Odontomachus troglodytes (Santschi): a behavioral intermediate stage in the evolution of social food exchange in ants. An Biol 17, 53–61.

23. Lois-Milevicich J, Schilman PE, Josens R. 2021 Viscosity as a key factor in decision making of nectar feeding ants. J Insect Physiol 128, 104164. (doi:10.1016/j.jinsphys.2020.104164)

24. Bonser R, Wright PJ, Bament S, Chukwu UO. 1998 Optimal patch use by foraging workers of Lasius fuliginosus, L. niger and Myrmica ruginodis. Ecol Entomol 23, 15–21. (doi:10.1046/J.1365-2311.1998.00103.X)

25. Dejean A, Solano PJ, Ayroles J, Corbara B, Orivel J. 2005 Arboreal ants build traps to capture prey. Nature 434, 973–973. (doi:10.1038/434973a)

26. Detrain C, Prieur J. 2014 Sensitivity and feeding efficiency of the black garden ant Lasius niger to sugar resources. J Insect Physiol 64, 74–80. (doi:10.1016/J.JINSPHYS.2014.03.010)

27. Falibene A, de Figueiredo Gontijo A, Josens R. 2009 Sucking pump activity in feeding behaviour regulation in carpenter ants. J Insect Physiol 55, 518–524. (doi:10.1016/J.JINSPHYS.2009.01.015)

28. Fujioka H, Okada Y. 2019 Liquid exchange via stomodeal trophallaxis in the ponerine ant Diacamma sp. from Japan. J Ethol 37, 371–375. (doi:10.1007/s10164-019-00602-9)

29. Blüthgen N, Menzel F, Blüthgen N. 2006 Measuring specialization in species interaction networks. BMC Ecol 6, 9. (doi:https://doi.org/10.1186/1472-6785-6-9)

30. VanGinkel CG, Gayton S. 1996 The biodegradability and nontoxicity of carboxymethyl cellulose (DS 0.7) and intermediates. Environ Toxicol Chem 15, 270–274. (doi:10.1002/ETC.5620150307)

31. Osswald T, Rudolph N. 2014 Polymer Rheology. In Polymer Rheology, pp. I–XI. München: Carl Hanser Verlag GmbH &amp; Co. KG. (doi:10.3139/9781569905234.fm)

32. Rhee BO, Lee S. 1999 Evaluation on Accuracy of the Rheological Data of PIM Feedstocks. Journal of the Japan Society of Powder and Powder Metallurgy 46, 830–836. (doi:10.2497/jjspm.46.830)

33. Emmerich A. 1994 Density data for sucrose solutions. Contribution to the application of the new values adopted by ICUMSA. Zuckerindustrie (Germany) 119, 120–123.

34. de Gennes P-G, Brochard-Wyart F, Quéré D. 2004 Capillarity and Wetting Phenomena. New York, NY: Springer New York. (doi:10.1007/978-0-387-21656-0)

35. Josens RB, Farina WM, Roces F. 1998 Nectar feeding by the ant Camponotus mus: intake rate and crop filling as a function of sucrose concentration. J Insect Physiol 44, 579–585. (doi:10.1016/S0022-1910(98)00053-5)

36. Yamanaka O, Takeuchi R. 2018 UMATracker: An intuitive image-based tracking platform. Journal of Experimental Biology 221. (doi:10.1242/JEB.182469/VIDEO-3)

37. Virtanen Pet al. 2020 SciPy 1.0: fundamental algorithms for scientific computing in Python. Nat Methods 17, 261–272. (doi:10.1038/s41592-019-0686-2)

38. Jean Leonard Poiseuille. 1844 Recherches experimentales sur le mouvement des liquides dans les tubes de tres-petits diametres. Imprimerie Royale. See https://books.google.co.jp/books?id=uBN1Q-IRzTMC{.

39. Ávila Núñez JL, Naya M, Calcagno-Pissarelli MP, Otero LD. 2011 Behaviour of Odontomachus chelifer (Latreille) (Formicidae: Ponerinae) Feeding on Sugary Liquids. J Insect Behav 24, 220–229. (doi:10.1007/S10905-010-9249-1/FIGURES/4)

40. Jandt J, Larson HK, Tellez P, McGlynn TP. 2013 To drink or grasp? How bullet ants (Paraponera clavata) differentiate between sugars and proteins in liquids. Naturwissenschaften 100, 1109–1114. (doi:10.1007/S00114-013-1109-3/FIGURES/2)

41. Borrell BJ. 2006 Mechanics of nectar feeding in the orchid bee Euglossa imperialis: pressure, viscosity and flow. Journal of Experimental Biology 209, 4901–4907. (doi:10.1242/JEB.02593)

42. Kim W, Gilet T, Bush JWM. 2011 Optimal concentrations in nectar feeding. Proc Natl Acad Sci U S A 108, 16618–16621. (doi:10.1073/PNAS.1108642108)

43. Nicolson SW, de Veer L, Köhler A, Pirk CWW. 2013 Honeybees prefer warmer nectar and less viscous nectar, regardless of sugar concentration. Proceedings of the Royal Society B: Biological Sciences 280. (doi:10.1098/RSPB.2013.1597)

44. Dejean A, le Breton J, Suzzoni JP, Orivel J, Saux-Moreau C. 2005 Influence of interspecific competition on the recruitment behavior and liquid food transport in the tramp ant species Pheidole megacephala. Naturwissenschaften 92, 324–327. (doi:10.1007/S00114-005-0632-2/FIGURES/2)

45. Buffin A, Mailleux AC, Detrain C, Deneubourg JL. 2011 Trophallaxis in Lasius niger: A variable frequency and constant duration for three food types. Insectes Soc 58, 177–183. (doi:10.1007/S00040-010-0133-Y/TABLES/2)

46. Greenwald E, Segre E, Feinerman O. 2015 Ant trophallactic networks: simultaneous measurement of interaction patterns and food dissemination. Sci Rep 5, 12496. (doi:10.1038/srep12496)

47. Bles O, Deneubourg J-L. Nicolis SC. 2018 Food dissemination in ants: Robustness of the trophallactic network against resource quality. Journal of Experimental Biology 221. (doi:10.1242/jeb.192492)

48. Greenwald EE, Baltiansky L, Feinerman O. 2018 Individual crop loads provide local control for collective food intake in ant colonies. Elife 7, e31730. (doi:10.7554/eLife.31730)

49. Padday JF, Pitt AR, Pashley RM. 1975 Menisci at a free liquid surface: surface tension from the maximum pull on a rod. Journal of the Chemical Society, Faraday Transactions 1: Physical Chemistry in Condensed Phases 71, 1919. (doi:10.1039/f19757101919)

50. Leboeuf AC et al. 2016 Oral transfer of chemical cues, growth proteins and hormones in social insects. Elife 5. (doi:10.7554/eLife.20375)

51. Hakala SM, Meurville MP, Stumpe M, Leboeuf AC. 2021 Biomarkers in a socially exchanged fluid reflect colony maturity, behavior, and distributed metabolism. Elife 10. (doi:10.7554/ELIFE.74005)

52. Hakala SMet al. 2022 Socially transferred materials: why and how to study them. Trends Ecol Evol (doi:10.1016/j.tree.2022.11.010)

