## Supplemental material for "*Diacamma* ants adjust liquid foraging strategies in response to biophysical constraints"

Supp. Table 1. Weight of each concentration of sugar water per 1 μL

| Sugar concentration % (w/w) | Weight (mg/μL) |
| --- | --- |
| 10 | 1.022 ± 0.005 |
| 20 | 1.0631 ± 0.008 |
| 30 | 1.1132 ± 0.003 |
| 40 | 1.1374 ± 0.006 |
| 50 | 1.1975 ± 0.010 |
| 60 | 1.2391 ± 0.011 |
| 10CMC | 1.0284 ± 0.005 |

Supp. Table 2. Viscosity.

CMC is a non-toxic inert viscosity modifier. 10CMC and 30CMC are 10% sugar solution with CMC 0.25% and 30% sugar solution with CMC 0.25% w/w.

| Sugar concentration % (w/w) | Viscosity (mPa•sec) |
| --- | --- |
| 10 | 1.158 ± 0.002 |
| 20 | 1.656 ± 0.002 |
| 30 | 2.641 ± 0.003 |
| 40 | 5.308 ± 0.005 |
| 50 | 11.919 ± 0.009 |
| 60 | 47.34 ± 0.004 |
| 10CMC | 8.97 ± 0.002 |
| 30CMC | 24.6 ± 0.2 |

Supp. Table 3. Drinking speed of each concentration of sugar water.

Drinking speed is the slope (μL/sec) of the linear regression of crop load and corresponding feeding time. Multiple comparison test was conducted to compare slopes using the non-parametric Wilcox test with Bonferroni correction (p< 0.05).

| Sugar concentration % (w/w) | Intake rate (μL/sec) | Intercept | Residual standard error | Wilcox test |
| --- | --- | --- | --- | --- |
| 10 | 0.0023 | 0.28 | 0.28 | a |
| 20 | 0.0032 | -0.42 | 0.43 | b |
| 30 | 0.0019 | 0.16 | 0.37 | abc |
| 40 | 0.0013 | 0.052 | 0.21 | c |
| 50 | 0.00089 | 0.062 | 0.17 | c |
| 60 | 0.00024 | 0.15 | 0.15 | d |
| 10CMC | 0.0011 | 0.13 | 0.2 | bc |

Supp. Table 4. The success rate of mandibular grabbing transportation to the nest. Ten ants’ return trips were followed from each of the three colonies.

| Sugar concentration % (w/w) | success | lost | colony |
| --- | --- | --- | --- |
| 10 | 30 | 0 | N = 3 |
| 50 | 30 | 0 | N = 3 |

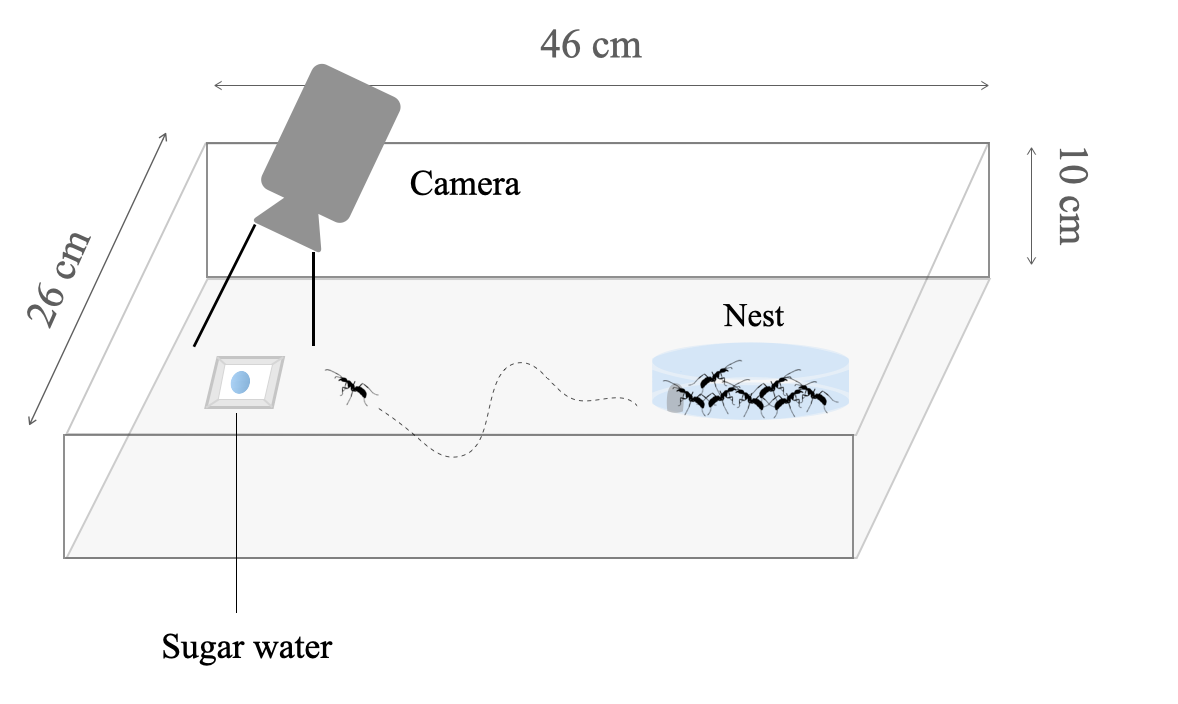
Supp. Figure 1. Experimental arena design.

An artificial nest (diameter = 9 cm) was placed in a larger arena container (48* 26* 10 cm). The nest was covered with a red film to darken inside of the nest. The distance between the nest and the food was approximately 30 cm. Fluon was applied to the walls of the foraging arena to prevent ants from escaping.

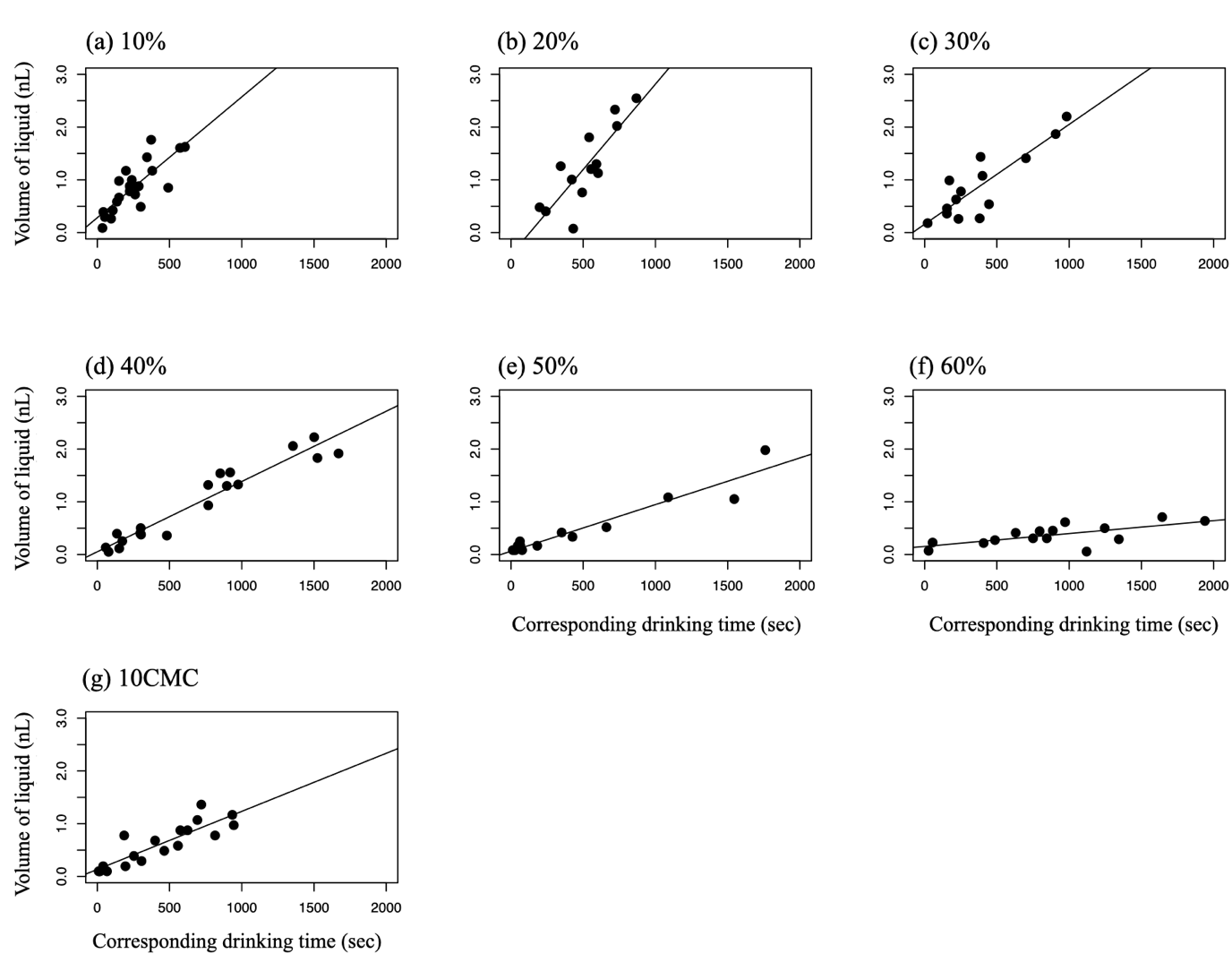
Supp. Figure 2. Drinking speed, plotting crop load (μL) and corresponding drinking time (sec).

(a-f) Each column indicates different sugar concentrations from 10% to 60% and (g) 10% sugar water with 0.25% CMC (10CMC). Linear regression models are shown in Supp. Table 3.

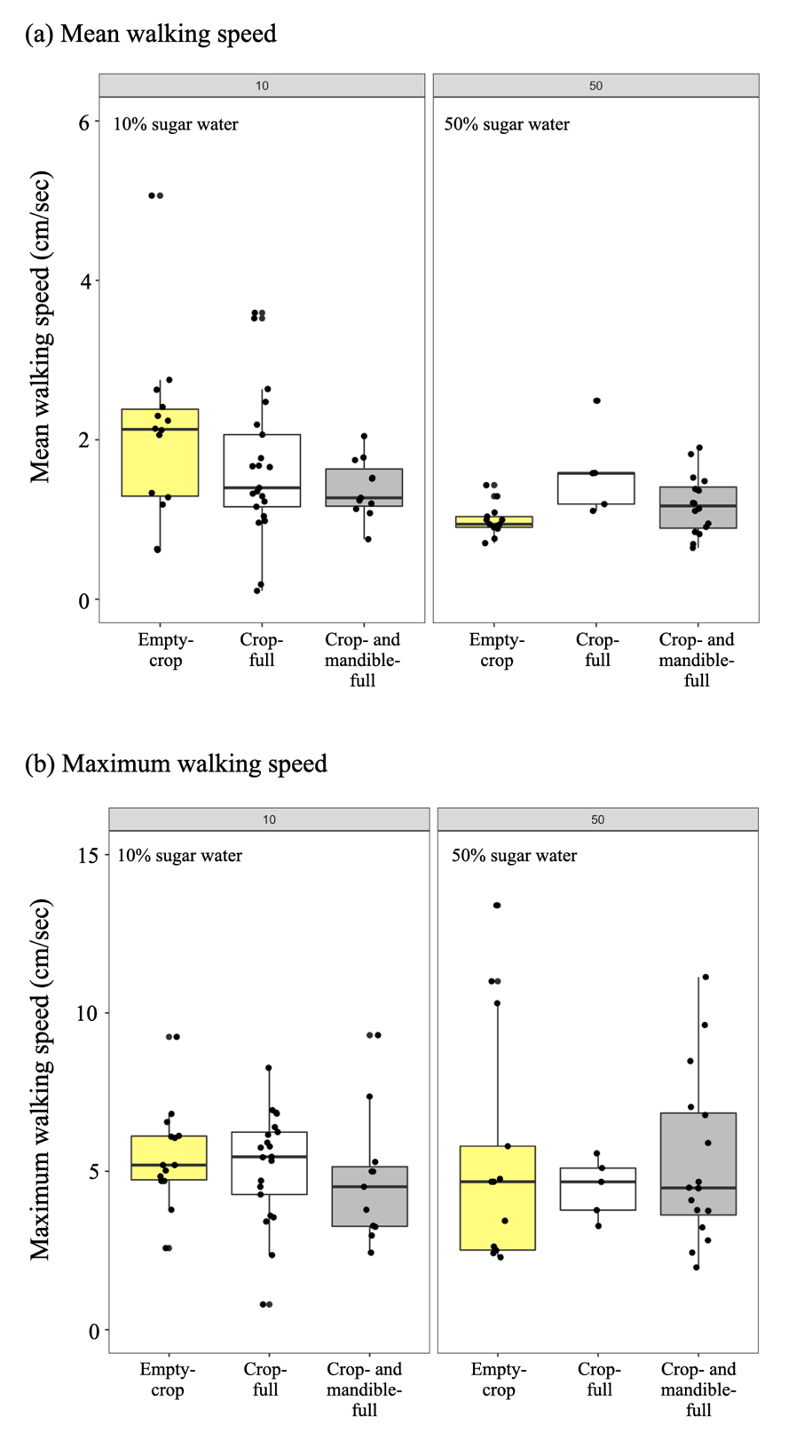

Supp. Figure 3. Walking speed.

Mean walking speed (a) and maximum walking speed (b) of ants across three conditions. There were no significant differences among the three categories (Tukey-Kramer test, p < 0.05).
